# Western Kenyan *Anopheles gambiae s.s.* showing intense permethrin resistance harbor distinct microbiota

**DOI:** 10.1101/2020.11.12.378760

**Authors:** Diana Omoke, Mathew Kipsum, Samson Otieno, Edward Esalimba, Mili Sheth, Audrey Lenhart, Ezekiel Mugendi Njeru, Eric Ochomo, Nsa Dada

## Abstract

**Background:** Insecticide resistance poses a growing challenge to malaria vector control in Kenya and around the world. Following evidence of associations between the mosquito microbiota and insecticide resistance, we comparatively characterized the microbiota of *An. gambiae s.s*. from Tulukuyi village, Bungoma, Kenya, with differing permethrin resistance profiles.

**Methods:** Using the CDC bottle bioassay, 133 2-3 day-old, virgin, non-blood fed female F_1_ progeny of field-caught *An. gambiae s.s*. were exposed to five times (107.5μg/ml) the discriminating dose of permethrin. Post bioassay, 50 resistant and 50 susceptible mosquitoes were subsequently screened for *kdr* East and West mutations, and individually processed for microbial analysis using high throughput sequencing targeting the universal bacterial and archaeal 16S rRNA gene.

**Results:** 47% of the samples tested (n=133) were resistant, and of the 100 selected for further processing, 99% were positive for *kdr* East and 1% for *kdr* West. Overall, 84 bacterial taxa were detected across all mosquito samples, with 36 of these shared between resistant and susceptible mosquitoes. A total of 20 were unique to the resistant mosquitoes and 28 were unique to the susceptible mosquitoes. There were significant differences in bacterial composition between resistant and susceptible individuals (F=2.33, P=0.001), with presence of *Sphingobacterium, Lysinibacillus* and *Streptococcus* (all known pyrethroid-degrading taxa), and the radiotolerant *Rubrobacter*, being significantly associated with resistant mosquitoes. On the other hand, the presence of *Myxococcus*, was significantly associated with susceptible mosquitoes.

**Conclusion:** This is the first report of distinct microbiota in *An. gambiae s.s*. associated with intense pyrethroid resistance. The findings highlight differentially abundant bacterial taxa between resistant and susceptible mosquitoes, and further suggest a microbe-mediated mechanism of insecticide resistance in mosquitoes. Our results also indicate fixation of the *kdr* East mutation in this mosquito population, precluding further analysis of its associations with the mosquito microbiota, but presenting the hypothesis that any microbe-mediated mechanism of insecticide resistance would be likely of a metabolic nature. Overall, this study lays initial groundwork for understanding microbe-mediated mechanisms of insecticide resistance in African malaria vectors, and potentially identifying novel microbial markers of insecticide resistance that could supplement existing vector surveillance tools.

## Background

Malaria remains an important global health problem, with 92% of all deaths occurring in Africa [1]. In Kenya, more than 70% of the population is at risk of the disease, with children aged ≤ 5 years and pregnant women being the most vulnerable to infection [2]. The use of indoor residual spraying (IRS), long-lasting insecticidal nets (LLINs) and other interventions have led to measurable improvements in preventing malaria [3]. Continued reliance on insecticide-based interventions has also resulted in widespread insecticide resistance in malaria vectors, thus threatening malaria control efforts [4, 5]. This is the case in western Kenya, where malaria vector control is increasingly being threatened by insecticide resistance due to selection pressure imposed by continued exposure to insecticides [4, 6].

Although insecticide resistance is increasingly prevalent [7], its underlying mechanisms are not fully understood. So far, four principal mechanisms of insecticide resistance have been described in mosquitoes, including: (i) metabolic resistance due to elevated activity of detoxification enzymes, (ii) target-site resistance due to genetic alterations at insecticide binding sites, (iii) cuticle modifications that prevent or reduce insecticide penetration, and (iv) behavioral changes resulting in avoidance of, or reduced contact with, insecticides [8]. Recent studies suggest that the mosquito microbiota may provide a fifth mechanism contributing to insecticide resistance [9, 10]. Focusing largely on *Anopheles albimanus* across different geographical locations including Peru [9] and Guatemala [10], these studies have identified significant alterations of the mosquito microbiota associated with insecticide resistance, with enrichment of insecticide-degrading bacteria and enzymes in resistant mosquitoes [9].

The mosquito microbiota has been shown to affect mosquito physiology [11]. These microbes, which are predominantly acquired during the aquatic life stage from aquatic habitats, colonize mosquito tissues including the gut, reproductive tracts, exoskeleton, and hemocoel [12, 13]. Some of these microbes are beneficial to mosquitoes through their role in nutrient provisioning, immunity and development, and subsequent contributions to mosquito fitness [12]. They also help provide protection against pathogens by modifying the host’s immune system or by synthesizing specific toxins [12]. The mosquito microbiota can influence and/or be influenced by several mosquito-related factors including mosquito species, developmental stage, genetics, and sex [11]. In malaria vectors, the microbiota play important roles in malaria parasite development, survival, and sporozoite prevalence, thus modulating vector competence [14–17].

Recent studies on the effects of insecticide exposure on microbes associated with malaria vectors and their habitats have so far focused on *An. stephensi, An. albimanus* and *An. arabiensis* [9, 10, 18, 19]. The microbiota of *An. gambiae s.s.* has, however, largely been unexplored in relation to insecticide resistance. Of particular importance is pyrethroid resistance—a major concern in Kenya, where this class of insecticide is predominantly used in LLINs and IRS [6, 20, 21]. To address this research gap, this study characterized and compared microbiota between pyrethroid resistant and susceptible *An. gambiae s.s.* from an area with intense pyrethroid resistance in Western Kenya. Mosquitoes were also screened for gene mutations that mediate knockdown resistance (*kdr*) to pyrethroids, in order to characterize any associations between the mosquito microbiota and *kdr* genotype. We discuss these findings on *An. gambiae s.s.*, and highlight their implications for insecticide resistance monitoring and management.

## Methods

### Mosquito collections

Mosquito collections were conducted in April and May 2018 in Tulukuyi village located in Bungoma County 0.56°N 34.56°E 1427m ASL (Figure 1). Previous studies conducted by Ochomo, *et al.* [20] indicated that *An. gambiae s.s.* was the most predominant species in Bungoma and had high resistance levels to pyrethroids. Sampling was performed by aspiration of blood fed and gravid mosquitoes from 39 houses. Mosquitoes were placed in labelled paper cups with information identifying the collection date and collection site. A piece of cotton wool soaked in 10% w/v sugar solution was placed on top of the netting material covering the paper cup to sustain the collected mosquitoes. The paper cups were then placed in a cool box and transported to the laboratory.

**Figure 1:**
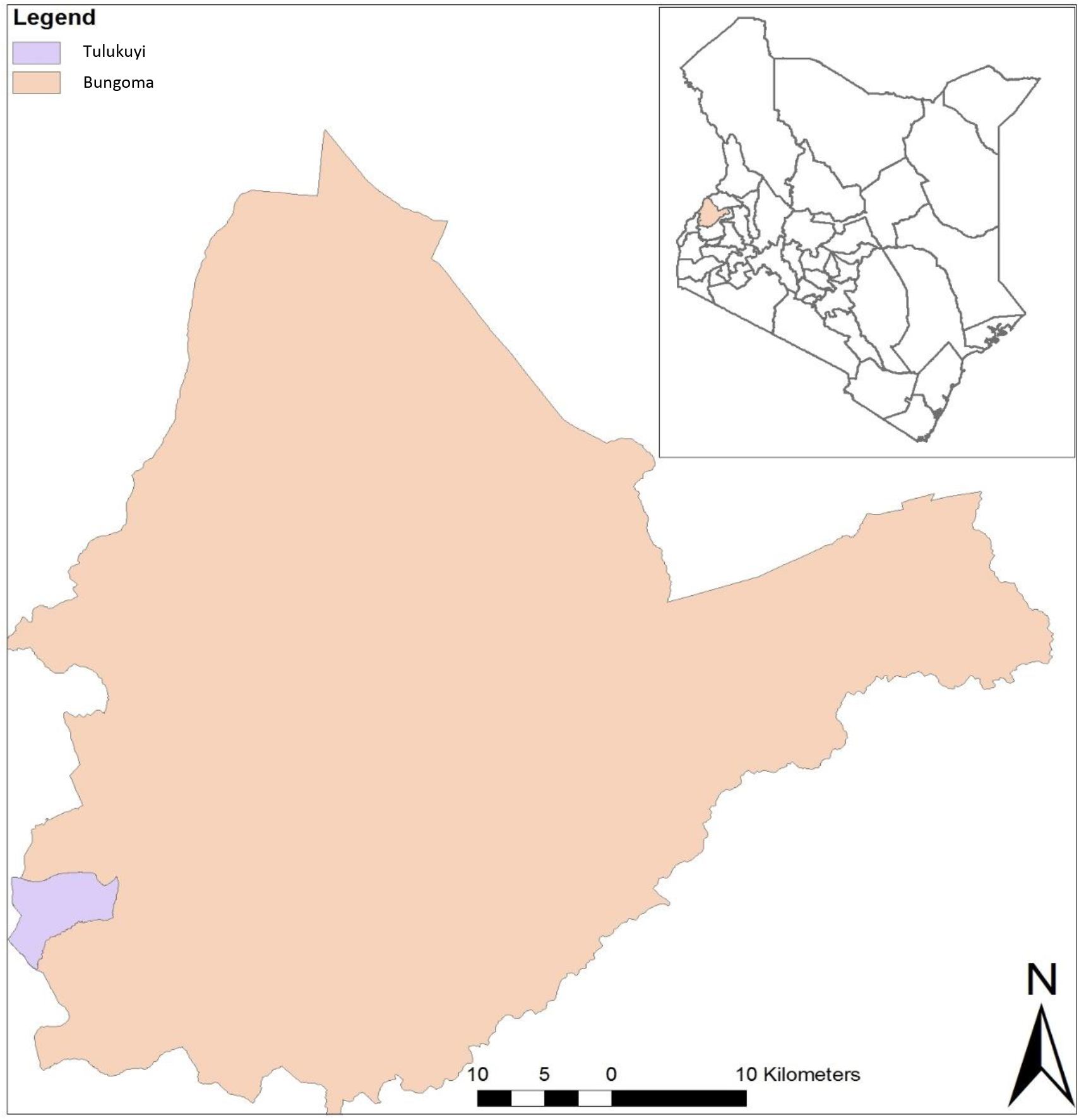
Map of Kenya (Right) showing Bungoma County (in expanded view) where adult mosquito collections were conducted. Adult female *Anopheles gambiae s.s*. were collected from Tulukuyi village and F_1_ progeny resulting from these mosquitoes were analyzed.

### Generation of F_1_ progeny from field-collected mosquitoes

Prior to species identification, forced oviposition was used to generate isofemale F_1_ progeny from field collected blood-fed and/or gravid female mosquitoes. Individual mosquitoes were placed in separate 50ml falcon tubes containing damp cotton wool topped with filter paper for egg laying. Following egg laying, each adult female was transferred into individual 1.5ml Eppendorf tubes for molecular species identification (described below). F_1_ eggs from each isofemale were removed and placed into separate clean larval trays containing distilled water for hatching, while the parents underwent species identification. Following species identification, all larvae from isofemales identified as *An. gambiae s. s.* were pooled, approximately 200 larvae per tray measuring 46cm by 35cm by 5cm and reared together. Larvae were fed a combination of ground TetraMin Baby^®^ fish food containing Brewer’s yeast (Spectrum Brands, Inc., WI), and Koi’s choice premium fish food (Foster & Smith, Inc. Rhinelander, WI) at a ratio of 1:2 for *An. gambiae* until pupation. Using a dissecting microscope (Nikon C-PS, model no. 1071990), male and female pupae were separated within 24 hours of pupation in order to obtain virgin adult females. Female pupae were subsequently placed into cages for adult eclosion, while the males were euthanized and discarded. The resulting F_1_ adult females (~377) were sustained on cotton balls soaked in 10% sugar solution for 2-3 days prior to insecticide susceptibility bioassays. All mosquito handling and rearing were conducted in the insectary at the Kenya Medical Research Institute - Center for Global Health Research (KEMRI-CGHR), Kisian, Kisumu, under the following conditions: temperature of 27 ± 2 °C, relative humidity of 80 ± 10 %, and photoperiod of 12:12 light: dark cycle.

### Molecular identification of mosquito species

Using the ethanol precipitation method described by Collins*, et al.* [22], genomic DNA was extracted from whole individual field-collected female mosquitoes that were used to generate the F_1_ progeny. 2 μl of DNA from each individual, along with known *An. gambiae s.s.* DNA as positive control, were used as templates for the PCR reaction [23]. The reactions were performed using BIO-RAD thermal cycler model T100 under the following conditions: 95°C for 5 min followed by 95°C for 30 sec, 56°C for 30 sec and 72°C for 30 sec for 30 cycles, with a final extension at 72°C for 5 min. Amplicons (~ 390bp for *An. gambiae s.s.*) were resolved by ethidium bromide-stained agarose gel (2%) electrophoresis.

### Permethrin resistance phenotyping

A total of 133 F_1_ virgin, non-blood fed adult females aged 2-3 days were tested for permethrin resistance following the Centers for Disease Control and Prevention (CDC) guidelines for evaluating insecticide resistance [24]. The control bottle was coated with 1ml of acetone while the four test bottles were coated with 1ml of permethrin stock solution prepared with acetone, at a final concentration of 107.5μg/ml (5X the dose for discriminating permethrin resistance in *Anopheles*). *Anopheles gambiae* Kisumu susceptible strain of the same age and physiological status were used to confirm the viability of the prepared bottles—all mosquitoes in the insecticide treated bottles died, while those in the acetone-treated bottles survived. Using the F_1_ progeny, the bioassays were conducted for 30 minutes at the end of which permethrin resistance was recorded. Mosquitoes that were alive after the bioassay were categorized as resistant and subsequently killed by freezing, while those that were dead or moribund were categorized as susceptible. Phenotyped mosquitoes were immediately placed in 1.5ml Eppendorf tubes with unique identification codes and preserved at −20°C for subsequent molecular processing.

### Genomic DNA isolation and molecular processing

### DNA isolation and purification

250 μl of 70% ethanol was added to each tube of individual F_1_ mosquitoes and vortexed at high speed for ~10 seconds to surface sterilize the mosquitoes. This was followed by a vigorous rinse, then a gentle rinse with 250 μl of nuclease free water each. Genomic DNA from the whole mosquito was isolated and purified using the MasterPure™ Gram Positive DNA Purification Kit following the manufacturer’s instructions (Epicentre Biotechnologies, Madison, USA). During DNA extraction, four blank controls (containing all the reagents used sans mosquito) and two 1 g soil samples from Kisian (as a distinct source of microbes) were also processed to catch any potential sample processing and cross contamination respectively. Purified DNA samples were stored at −20°C for subsequent analysis.

### Detection of *kdr*-East and *kdr*-West alleles

RT-PCR was used to detect the presence of both *kdr*-East and *kdr* - West alleles using DNA from F_1_ mosquitoes. Following the protocol by Bass*, et al.* [25], samples were processed using the MxPro-Mx3005P software ‘Allele Discrimination-SNP’s’ program with 1.5 μl of DNA as template. PCR was carried out under the following cycling conditions for *kdr*-East: 95°C for 10 min then 40 cycles of 95°C for 10secs and 60°C for 45 sec. For *kdr*-West, the cycling conditions were 95°C for 10 min followed by 40 cycles of 92°C for 15 sec and 60°C for 60 sec.

### Library preparation and 16S rRNA gene amplicon sequencing

Using the 341f (**TCGTCGGCAGCGTCAGATGTGTATAAGAGACAGCCTACGGGNGGCWGCAG**) and 805r (**GTCTCGTGGGCTCGGAGATGTGTATAAGAGACAG** GACTACHVGGGTATCTAATCC) primers [26] with Illumina ® (San Diego, CA USA) overhang (bold typeface), the V3-V4 region of the universal bacterial and archaeal 16S rRNA gene was amplified using genomic DNA from F_1_ mosquitoes. Four no-template controls (PCR grade water), along with the six controls from the DNA extraction step—two cross-contamination controls (soil samples), and four blanks—were also processed. The PCR reaction mixture (25 μl total volume) comprised of 10 μl of 2X KAPAHiFi HotStart Mix (Roche, Switzerland), 5 μM each of 341f and 805r primers and 5 μl of DNA template which was ≥20 ng/μl. Reactions were conducted using the BIO-RAD T100 thermal cycler with the following cycling conditions: 95°C for 3 min for initial denaturation, followed by 25 cycles of 95 °C for 30 sec, 55 °C for 30 sec, 72 °C for 30 sec and extension at 72 °C for 5 min. The resulting amplicons of ~ 460 bps were purified using Agencourt AMPure XP beads (Beckman Coulter Inc., Indianapolis, IN, USA) at 0.87 x sample volume and eluted in 45 μL TE buffer. The purified amplicons including those from blank and cross contamination controls were submitted to the Biotechnology Core Facility at the US Centers for Disease Control and Prevention, Atlanta for library preparation and sequencing.

Sequencing libraries were obtained using index PCR. This comprised NEBNext Hig-Fidelity 2X PCR master mix New England Biolabs Inc., Ipswich, MA), index primers from Nextera XT Index kit v2 set A, B and D; (Illumina, San Diego, CA), and 10-300 ng of each 16S rRNA gene amplicon, along with all controls, as template. PCR thermal cycler conditions were set to: 98 °C for 30 sec, followed by 8 cycles of 98 °C for 10 sec, 55 °C and 65 °C for 30 sec each, followed by a final extension at 65 °C for 5 min. The resulting products were cleaned using Agencourt AMPure XP beads at 1.2 x volume of each library. These were subsequently analyzed for size and concentration, normalized and pooled at a final concentration of 2 nM. The pool was denatured using Illumina guidelines for loading onto flowcell for cluster generation, and sequencing was performed on an Illumina Miseq using Miseq 2×250 cycle paired-end sequencing kits. The resulting sequence reads were filtered for read quality, basecalled and demultiplexed using bcl2fastq (v2.19.1).

### Sequencing data quality control and generation of amplicon sequence variants (ASV) table

Resulting raw paired-end sequencing reads were demultiplexed and imported into the Quantitative Insights Into Microbial Ecology (QIIME) 2 pipeline v.2018.11 [27] for analysis. Primers and adapter sequences were removed using the QIIME2 cutadapt plugin v.2018.11.0 [28]. This was followed by quality filtering using the QIIME2 DADA2 plugin v.2018.11.0 to remove any sequencing errors, denoise and dereplicate paired-end sequences, filter out chimeras, and finally generate a frequency table of Amplicon Sequence Variants (ASVs, also referred hereafter as features) [29]. The quality filtering step was achieved using the denoise-paired command with the following parameters; `max_ee: 2, trunc_q: 2, trim_left_f: 10, trim_left_r: 10, n_reads_learn: 1000000` and all other parameters left as default. The resulting frequency table was subsequently filtered to remove features associated with the controls, and those with frequency <100 prior to downstream analysis. Following these steps, 36 susceptible and 39 resistant mosquitoes remained, and were used for downstream analysis. The raw sample sequencing reads generated from this project, including those from negative (blank) and cross contamination (soil samples) controls, along with sample metadata, have been deposited in the National Center for Biotechnology Information (NCBI), Sequence Read Archive under the BioProject PRJNA672031

### Microbial community diversity analysis

Alpha and beta diversity indices [30] were computed and compared between samples with differing resistance phenotypes. Shannon alpha diversity index, a quantitative measure of community richness and evenness, was computed using the q2-diversity plugin. To avoid introducing bias due to unequal sampling depth, prior to alpha diversity analysis, all samples were rarefied to a depth of 100 ASVs per sample (Suppl. 2), which was sufficient to capture the typical low microbiota diversity in individual mosquitoes. The Kruskal-Wallis test was used to compare Shannon diversity indices between insecticide resistant and susceptible samples with Benjamini-Hochberg false discovery rate (FDR) corrections.

Bray-Curtis dissimilarity beta diversity index, a quantitative measure used to determine compositional dissimilarity of features between samples, was also computed using the `q2-diversity` plugin. The Bray-Curtis dissimilarity matrices were computed using both rarefied (to a sampling depth of 100 ASVs per sample as described above) and unrarefied ASVs. Both resulted in significant differences between the microbiota of resistant and susceptible mosquitoes (Suppl. 3), thus, ordination outputs of only the latter are presented. Comparisons of the resulting distance matrices between resistant and susceptible samples were performed using permutational multivariate analysis of variance (PERMANOVA) at 999 permutations with Benjamini-Hochberg FDR corrections. Outputs were visualized using phyloseq package [31] in R [32]

### Taxonomic annotation of microbial features

QIIME2 v 2018.11 `q2-feature-classifier` plugin [33] was used for taxonomic annotation. The Naïve Bayes classifier [34] was pre-trained on 16S SILVA reference (99% identity) database v.128 [35]. Using the qiime `feature-classifier extract-reads` command, trimming was done to only target the V3-V4 region of the 16S rRNA gene (~ 425 bps length). The `qiime feature-table heatmap` plugin was subsequently used to visualize the resulting relative abundance of annotated ASVs across samples. The plugin’s metrics and clustering methods were set to `braycurtis` and `features` respectively.

### Testing for differentially abundant microbial features between permethrin resistant and susceptible mosquitoes

The linear discriminant analysis (LDA) effect size method (LEfSe) [36] was used to identify ASVs that were differentially abundant between resistant and susceptible mosquitoes. Annotated ASVs were converted into abundance tables and uploaded to LEfSe Galaxy v.1.0 (http://huttenhower.sph.harvard.edu/lefse/). With default parameters, an alpha value of 0.05 was used for both the factorial Kruskal-Wallis and pairwise Wilcoxon tests within LEfSe, and a threshold value of >2 was used on the resulting logarithmic LDA score to identify differentially abundant ASVs. The effect sizes of differentially abundant ASVs were visualized as bar plots.

The analysis of composition of microbiome method, ANCOM [37], was used to verify the results obtained from LEfSe. The ANCOM analysis was called using the `q-2 composition` plugin, with the transform and difference functions set to `log_transform` and `mean_difference`, respectively. All other parameters were set to default. The resulting statistic, W, and its default cut off was used to identify differentially abundant features between resistant and susceptible mosquitoes.

## Results

### Summary statistics of permethrin resistance phenotypes, *kdr* mutations and sequencing data

A total of 133 adult F_1_ female *An. gambiae s.s.* were tested for resistance to permethrin using 5X (107.5 μg/ml) the discriminating dose (21.5 μg/ml) of the insecticide, and 52.6% of the samples tested were found to be susceptible. One hundred of the screened samples (50 resistant and 50 susceptible) were subsequently processed for characterizing the microbiota and *kdr* allele frequencies. Of all 100 samples, 99% had the *kdr* east (*Vgsc_1014S*) mutation and the remaining one had the *kdr* west (*Vgsc_1014F*) gene mutation. This high frequency of *kdr* east mutation indicated fixation in the mosquito population and thus precluded further correlation analysis between microbial composition and *kdr* allele frequencies. Microbial community characterization of all 100 samples yielded 4,319,065 raw sequencing reads, in addition to 5,226 raw reads from blank and cross-contamination controls (Suppl. 1). Following sequencing data quality control and subsequent removal of features associated with controls and those with frequency < 100, 36 susceptible and 39 resistant samples remained and were used for downstream analysis

### Microbiota composition differed between permethrin resistant and susceptible *An. gambiae s.s*

Comparison of Bray-Curtis dissimilarity indices using PERMANOVA, showed significant differences in bacterial composition between permethrin resistant and susceptible mosquitoes (pseudo-F=2.33, *p*=0.001). This heterogeneity in microbial community structure associated with insecticide resistance status was further illustrated by principal coordinates analysis (PCoA), in which the microbiota of susceptible samples clustered closely together and away from those of primarily dispersed resistant samples (Figure 2).

**Figure 2:**
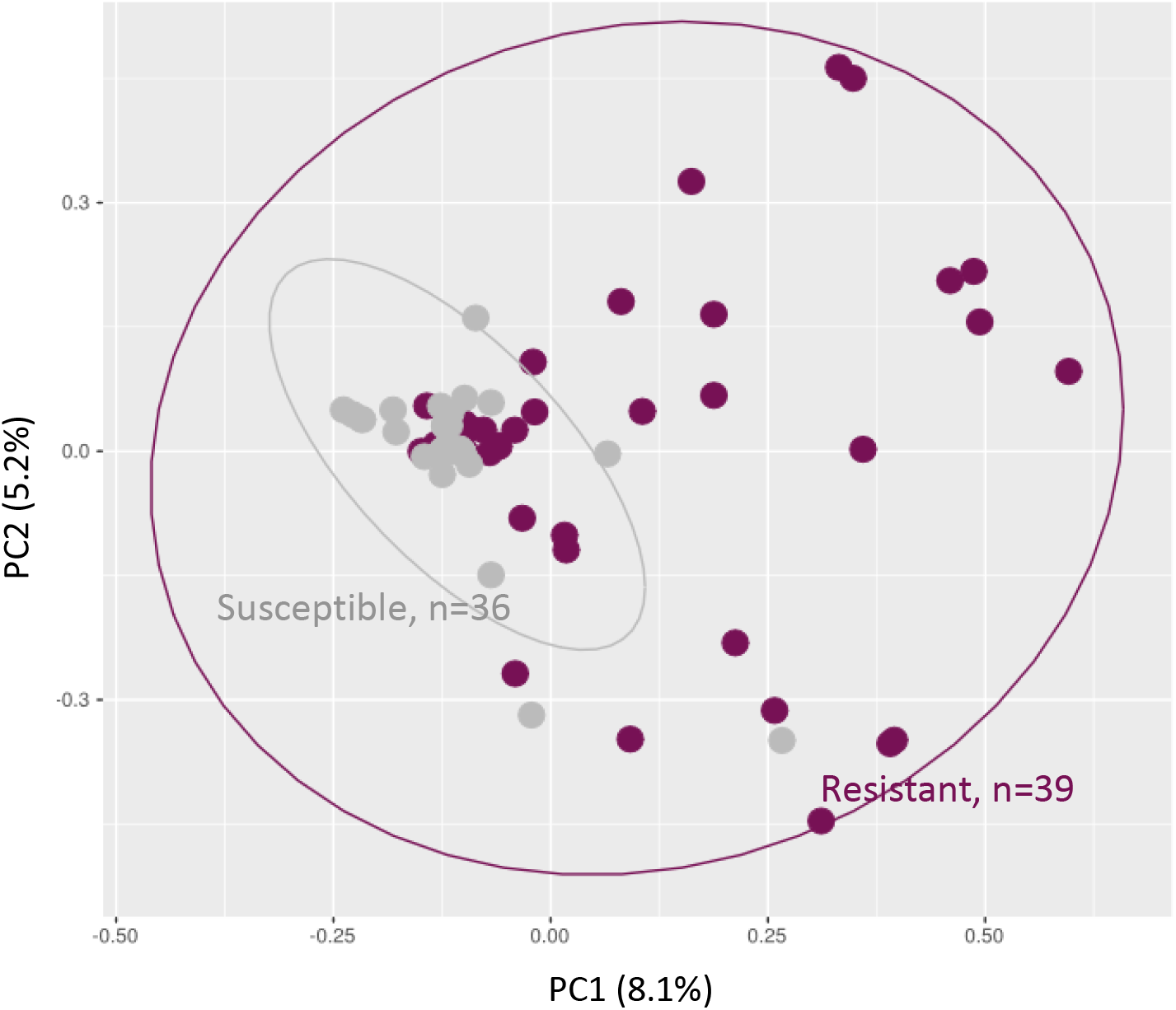
A Principal Coordinate analysis (PCoA) plot of Bray-Curtis distances between the microbiota of permethrin resistant and susceptible *An. gambiae s.s.* Each point on the plot represents the microbial composition of a single mosquito. The susceptible samples clustered closely together and away from the primarily dispersed resistant samples. The Bray-Curtis comparison using permutational multivariate analysis of variance (999 permutations) showed a significant difference in microbial composition between resistant and susceptible samples (pseudo-F=2.33, *p*=0.001).

Considering microbial diversity within each group, Kruskal-Wallis comparison showed no difference in Shannon diversity between the microbiota of permethrin resistant and susceptible mosquitoes (H = 0.45, *p* = 0.50) (Suppl. 4)

### *Anopheles gambiae s.s.* from Tulukuyi comprised sparse but diverse microbial taxa that differed by permethrin susceptibility status

Taxonomic annotation was performed to the genus level or to the lowest possible taxonomic rank. The relative frequencies of annotated bacterial taxa for each sample are presented in Figure 3. Overall, ASVs from *An. gambiae s.s*. microbiota were assigned to 84 bacterial taxa (Suppl. 5), and out of these, less than half (36 taxa) were shared between permethrin resistant and susceptible *An. gambiae s.s.*. There were 28 and 20 unique bacterial taxa in permethrin susceptible and resistant samples, respectively (Figure 3a, Suppl 4). At the genus level, a total of 66 bacterial genera were identified, 29 of which were shared between resistant and susceptible mosquitoes, while 21 and 16 were unique to permethrin susceptible and resistant mosquitoes respectively (Figure 3b, Suppl. 5).

**Figure 3:**
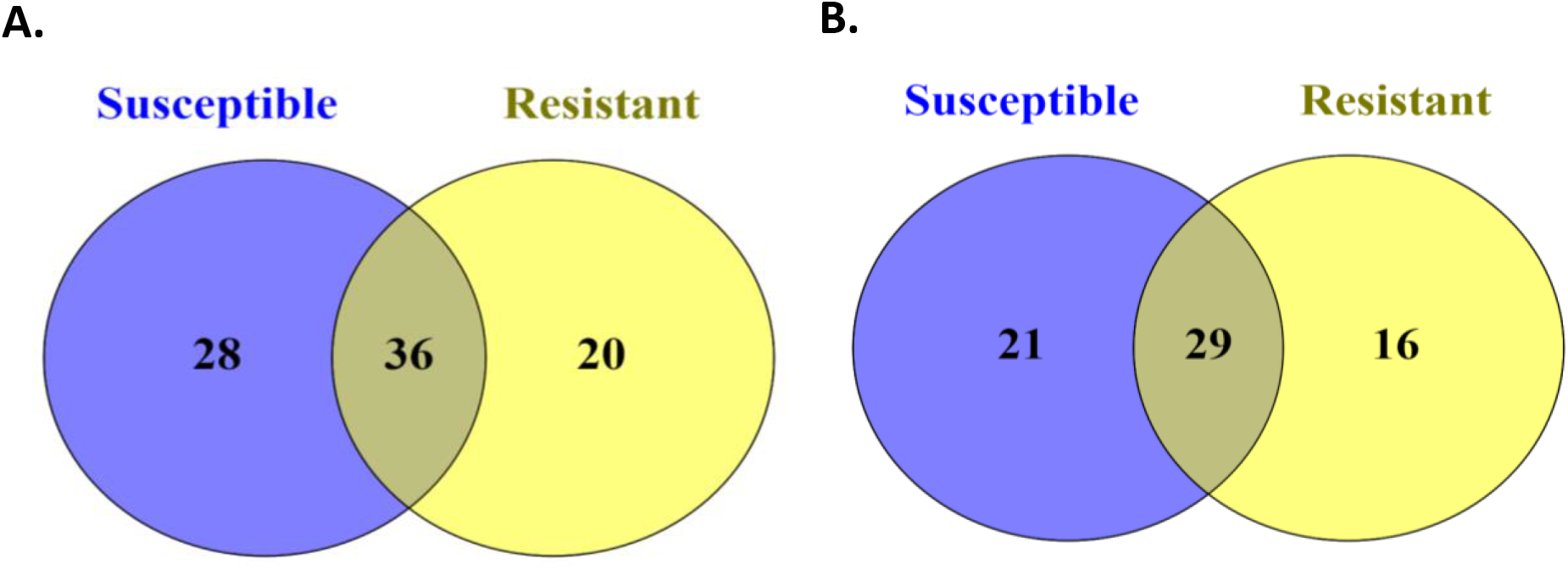
Venn diagrams showing number of bacterial taxa unique to or shared between 39 permethrin resistant and 36 susceptible mosquitoes. Panel A. shows number of bacterial taxa annotated to the genus or lowest possible taxonomic rank, and B. shows number at the genus level.

The most abundant bacterial taxa across all samples were those assigned to *Asaia* (38.33%), *Enterobacter* (7.25%), *Acinetobacter* (3.88%), *Klebsiella* (3.84%), an uncharacterized *Enterobacteriaceae* (3.30%), and *Lysinibacillus* (3.27%), together accounting for more than 55% of ASVs (Suppl. 5). A total of 16 genera were unique to resistant mosquito samples including *Lysinibacillus, Thorsellia, Streptococcus* and *Altererythrobacter*, among others (Suppl 4). The six most dominant genera among resistant mosquitoes were *Lysinibacillus* (13.97%), *Pseudomonas* (11.95%), *Acinetobacter* (8.54%), *Thorsellia* (6.49%), *Asaia* (4.23%) and *Bacillus* (4.08%). On the other hand, 21 genera were only found in the susceptible mosquito samples including *Marmoricola, Roseomonas, Dyadobacter, Lactococcus,* and *Myxococcus*, among others. Among susceptible mosquitoes, *Asaia* was the most dominant, with a relative abundance of 48.76% followed by *Enterobacter* (9.23%), *Klebsiella* (4.41%), *Enterococcus* (3.63%) and *Acinetobacter* (2.45%).

A few resistant and susceptible individuals had highly diverse microbiota, with ASVs assigned to between 14 and 37 bacterial taxa (Figure 4 and Suppl.4). The sample with the highest bacterial diversity was a permethrin susceptible mosquito. Notably, some bacterial taxa were detected more frequently in resistant compared to susceptible mosquitoes. These included the genus *Rubrobacter* which was detected at low abundance in 20 of the 39 permethrin resistant samples and only in two susceptible mosquito samples, also at low levels of abundance. Similarly, ASVs assigned to unclassified *Rhodospirillales* (JG37-AG-20) and unclassified *Obscuribacteriales* were detected in 18 and 9 resistant mosquitoes, respectively, but only in 1 and 4 susceptible mosquitoes, respectively. ASVs assigned to the genera *Streptococcus*, *Thermomonas*, *Sphingobacterium*, *Ornithinimicrobium* and *Lysinibacillus* were detected in more permethrin resistant samples compared to the susceptible samples (Figure 4 and Suppl.4). On the other hand, ASVs annotated as unclassified *Enterobacteriaceae* were predominant in the susceptible mosquitoes and were detected in 10 of these samples in contrast to only 4 resistant samples.

**Figure 4.**
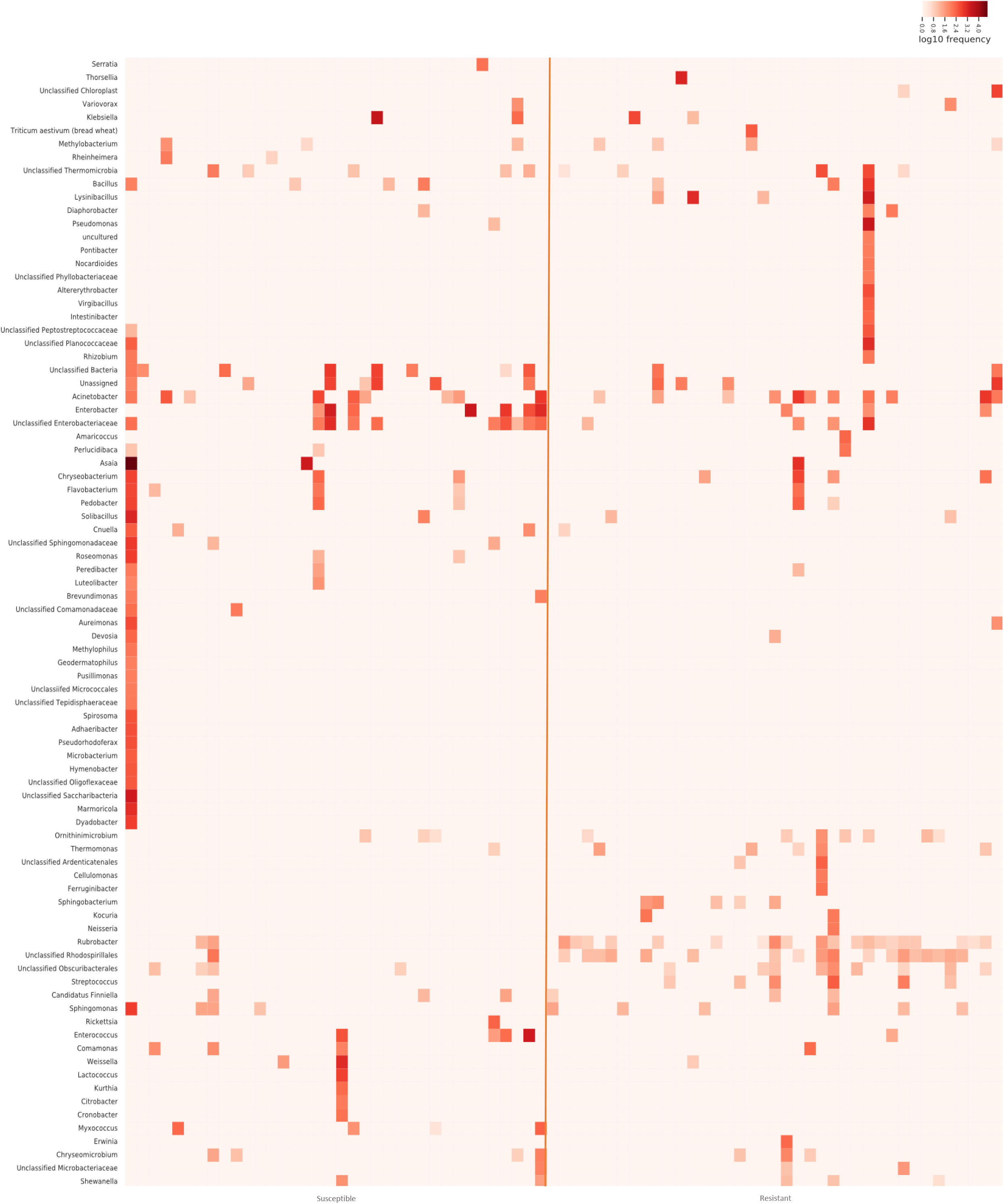
Heatmap showing frequency of annotated ASVs. Frequency of ASVs from the microbiota of individual permethrin resistant (n = 39) and susceptible (n = 36) *An. gambiae s. s.* from Tulukuyi. The annotation of ASVs was done to the genus level or lowest possible taxonomic level.

### Differentially abundant bacterial taxa between insecticide resistant and susceptible mosquitoes

Linear discriminant analysis (LEfSe) also revealed significant differences in microbiota composition between susceptible and resistant mosquitoes. Focusing on the genus level, four bacterial genera, *Sphingobacterium, Streptococcus, Lysinibacillus*, and *Rubrobacter*, and an uncultured bacterium were highlighted by LEfSe as more abundant in resistant mosquitoes (Figure 5). The first three genera were only detected in resistant mosquitoes, while *Rubrobacter* and the uncultured bacterium were at least three-fold more abundant in resistant compared to susceptible mosquitoes (Figure 5 and Suppl. 5). On the other hand, LEfSe identified only one bacterial genus, *Myxococcus,* as more abundant in the susceptible samples (Figure 5); this genus was not detected at all in the resistant samples (Suppl. 5). Although more bacterial genera were unique to either resistant or susceptible mosquitoes (Figure 3b and Suppl. 5), LEfSe highlighted those that were present in at least four individuals.

**Figure 5:**
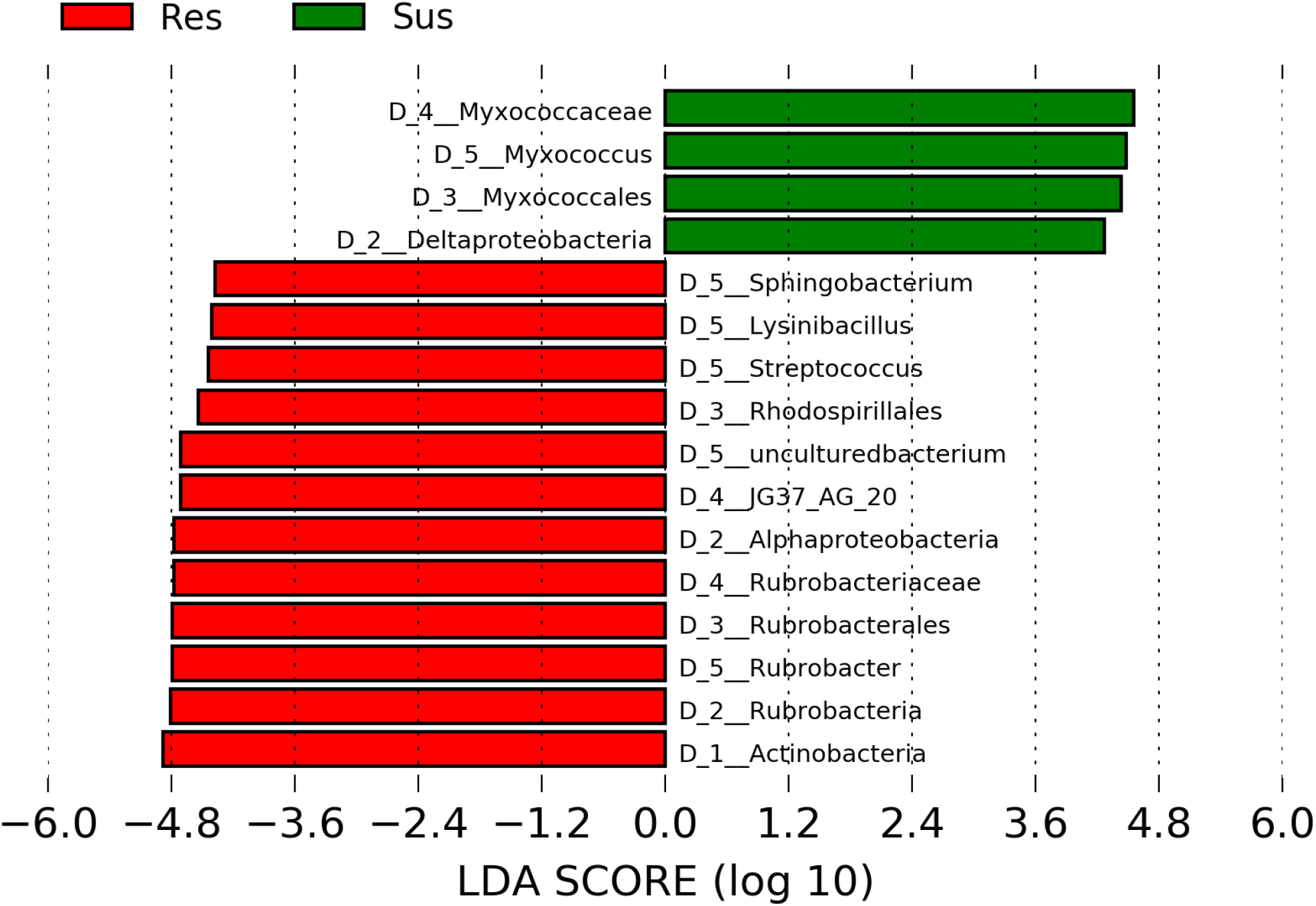
Differentially abundant bacterial genera between permethrin resistant and susceptible mosquitoes. The green and the red bars represent taxa which were significantly more abundant in the susceptible and resistant samples, respectively, at log 10 transformation. Taxonomic levels are designated as D_1_phylum, D_2_class, D_3_order, D_4_family, D_5_genus.

The ANCOM method further corroborated these results. Being more stringent, and not considering features that were unique to either sample category, it identified features assigned to the genus *Rubrobacter* (W= 51) and unclassified *Rhodospirillales* (JG37-AG-20) (W = 63) as significantly more abundant in resistant compared to susceptible samples (Suppl. 6).

## Discussion

Recently, studies of *An. stephensi, An. arabiensis* and *An. albimanus* have shown links between mosquito-associated microbiota and resistance to pyrethroids and organophosphates [9, 10, 18, 19]. In this study we comparatively characterized the microbiota between pyrethroid resistant and susceptible F_1_ progeny of field-derived *An. gambiae s.s*. Our results showed significant differences in microbiota composition between resistant and susceptible mosquitoes with enrichment of different bacterial taxa between resistant and susceptible mosquitoes. We detected intense resistance (at 5X the diagnostic dose) to permethrin, along with high frequency (99.14%) of the *kdr* east allele in the F_1_ progeny originating from Tulukuyi, Western Kenya. These findings corroborate earlier reports of high pyrethroid resistance in the same area [6, 20]. Multiple studies from western Kenya have indicated that the high intensity of insecticide resistance may be contributing to mosquito control failure [20, 21]. The high frequency of the *kdr* east allele suggests that the mutation is fixed in this mosquito population. Other studies conducted in western Kenya have also reported the presence of high *kdr east* allele frequencies which is attributed to the continued use of insecticide–based vector control methods [20, 38–40]. However, our results showed that the allele was fixed regardless of resistance phenotype, suggesting that additional mechanisms, such as the overexpression of detoxification enzymes (e.g. cytochrome P450s [41]), are more important than *kdr* in conferring the intense permethrin resistance detected in the population. The fixation of the *kdr* east mutation in the population also precluded further analysis of any associations between *kdr* alleles and the mosquito microbiota. Indeed, a recent study identified no links between the two [42]. We thus hypothesize that any microbe-mediated mechanism of insecticide resistance would be largely distinct from the mosquito host’s genetics, and likely of a metabolic nature.

Our results showed diverse bacterial taxa from individual *An. gambiae s.s.* samples, a majority of which have previously been identified in *Anopheles* and other mosquito genera including *Aedes aegypti* [43–46]. However, less than half of the detected microbial taxa were shared between permethrin resistant and susceptible mosquitoes, suggesting insecticide resistance-related physiological differences that favored different bacterial taxa.

Our results also showed significant differences in microbiota composition and structure between permethrin resistant and susceptible *An. gambiae s.s*. There is evidence that insecticide detoxifying microbes in agricultural insect pests contribute to insecticide resistance in their hosts [47, 48]. Recent studies on mosquitoes have also identified insecticide resistance- and/or exposure-driven alterations of the host microbiota. In particular, *Anopheles albimanus* microbiota differed by resistance to fenitrothion and was altered by exposure to different pyrethroids, and *Aedes aegypti* microbiota differed by resistance to lambda-cyhalothrin [9, 10, 49]. These findings suggest that insecticide resistance in mosquitoes favor the proliferation of certain bacterial taxa, possibly those that can degrade and metabolize insecticides. Recent studies [9, 10] identified known insecticide-metabolizing bacterial taxa in *Anopheles albimanus* that were exposed or resistant to insecticides. Huang*, et al.* [50] and Tang*, et al.* [51] documented that certain microorganisms (considered as potential candidates for bioremediation), including bacteria, degrade pesticides in the soil by breaking them down into smaller compounds, utilizing them as their source of nutrients and making them less toxic to the environment. Some of these microorganisms degrade pesticides to create conducive environments for their survival and not for nutritional requirements [51]. The different taxa present in the resistant vs susceptible mosquitoes, particularly those of resistant mosquitoes, is suggestive of this type of adaptation.

Despite significant differences in microbiota composition and structure (beta diversity), there was no significant difference in alpha (Shannon) diversity between the microbiota of resistant and susceptible mosquitoes. This is suggestive of a homeostatic-controlled number of microbial taxa across individual mosquitoes, with an insecticide resistance-associated perturbation of the type and relative abundance of specific microbial taxa. Mosquitoes used in this study were F_1_ progeny of wild adult females collected from the same location and reared under identical conditions. Except for their permethrin resistance status, which was determined at 2-3 days post adult eclosion, the mosquitoes had identical physiological characteristics. These identical rearing conditions and subsequent uniform physiological characteristics may explain the homogeneity in alpha diversity across samples. On the other hand, the differences in microbial composition associated with their permethrin resistance status provide further evidence of insecticide selection pressure on the mosquito microbiota. It is well known that a majority of the mosquito microbiota is obtained from mosquito aquatic habitats at the larval stage, and also from food sources as adults [13]. Newly emerged adults can also imbibe bacteria along with water from their larval habitats during eclosion or through transstadial transmission [52]. However, other factors such as mosquito physiological status [11, 52, 53] affect what microbes persist and colonize the mosquitoes following acquisition, and this could explain the insecticide resistance-associated differences in composition despite similar alpha diversity across all individual samples.

Differential abundance testing identified *Sphingobacterium*, *Lysinibacillus*, *Streptococcus* and *Rubrobacter* as significantly more abundant in resistant mosquitoes and *Myxococcus* as significantly more abundant in susceptible mosquitoes. The first three genera were only detected in resistant mosquitoes, while *Rubrobacter* was at least three-fold more abundant in resistant compared to susceptible mosquitoes. In a study conducted by Hu*, et al.* [54], *Lysinibacillus sphaericus* was identified as a microbe with the ability to degrade up to 83% of cyfluthrin (a pyrethroid) after 5 days of incubation by utilizing the insecticide as its source of carbon or nitrogen. In the current study, *Lysinibacillus* was only detected in resistant mosquitoes, likely as a result of its ability to utilize pyrethroids. Lozano and Dussán [55] also described the potential of *Lysinibacillus sphaericus* to be used in bioremediation of heavy metals. *Sphingobacterium* and *Streptococcus,* also only detected in resistant mosquitoes in this study, are bacterial genera known to degrade pyrethroid insecticides such as cypermethrin [56–58]. Bacteria belonging to the genera *Streptococcus* and *Rubrobacter* have been categorized as core microbiota of the digestive system of *Anopheles culicifacies* [59]. Although not documented for pyrethroid degradation or metabolism, *Rubrobacter* are known to be thermophilic and extremely resistant to UV thermal and gamma radiations [60]. Some other bacterial genera belonging to *Actinobacteria*, the phylum to which *Rubrobacter* belongs, have been associated with degradation of insecticides including pyrethroids [61, 62], and the overabundance of *Rubrobacter* in insecticide resistant mosquitoes could suggest their contribution to resistance. On the other hand, the genus *Myxococcus* was only detected in susceptible mosquitoes. This bacterial genus is known to be predatory on other bacteria [63], chitinase-producing [64], capable of producing various bioactive antifungal agents [65], and inhibitors of cellular respiration [66]. However, their association with mosquito physiology or insecticide susceptibility has not yet been described. Given what is known about this bacterial genus, it is possible that they could also be toxic to mosquitoes by directly inhibiting host’s cellular respiration and/or indirectly preying on other members of the mosquito microbiota that are necessary for host’s survival and or insecticide metabolism. Further studies are necessary to elucidate the role of *Myxococcus* and their secondary metabolites on mosquito physiology, including insecticide susceptibility.

In an aquatic microcosm, it has been demonstrated that insecticides, if used singly or in combination, can reduce microbial diversity and/or induce shifts in microbial community structure [67]. Recent studies have also demonstrated shifts in mosquito microbiota and larval water microbiota that were associated with insecticide exposure [10, 67]. This indicates that insecticide exposure shapes the microbial composition of mosquitoes and their habitats. This is likely due to the toxic effects of insecticides on some microbes, while at the same time favoring the proliferation of other tolerant microbes as described by Johnsen*, et al.* [68]. It is also possible that in addition to, or rather than selection pressure, the presence of specific insecticide-metabolizing microbes in mosquitoes induce resistance to insecticides and precludes colonization by other microbes. In *Aedes aegypti* it has been demonstrated that infections with certain microbes precludes colonization by others [69], and that microbial interactions within mosquitoes shape their microbial community [9, 70]. Further research on these microbial networks could shed more light on the role of the mosquito microbiota in insecticide resistance. [10, 67].

## Conclusion

In this study, we detected intense permethrin resistance in F_1_ progeny of field-collected *An. gambiae s.s.* from Tulukuyi, Bungoma, western Kenya. This was accompanied by a high frequency of (> 99%) of the *kdr* east allele, suggesting fixation in the population. We also show, for the first time, significant differences in microbiota composition between permethrin resistant and susceptible *An. gambiae s.s.* These findings corroborate results of previous research on other *Anopheles* species from different geographic locations. The abundance of *Rubrobacter*, *Lysinibacillus*, *Sphingobacterium* and *Streptococcus* were associated with resistant mosquitoes, while the abundance of *Myxococcus* was associated with susceptible mosquitoes. The enrichment of these specific bacterial taxa highlights the potential for discovering novel microbial markers of insecticide resistance that could complement existing insecticide resistance surveillance tools. With this increasing evidence of associations between mosquito microbiota and insecticide resistance, future work will evaluate the underlying microbial mechanisms of insecticide resistance.

## List of abbreviations

ASL: Above sea level
ASVs: Amplicon Sequence Variants
CDC: Centre for Disease Control
FDR: False discovery rate
IRS: Indoor residual spraying
Kdr: Knock Down Resistance gene
KEMRI-CGHR: Kenya Medical Research Institute-Center for Global Health Research
LDA: Linear Discriminant analysis
LEfSe: Linear discriminant analysis effect size
LLINs: Long Lasting Insecticidal Nets
PCoA: Principal Co-ordinates Analysis
PERMANOVA: Permutational multivariate analysis of variance
QIIME: Quantitative Insights Into Microbial Ecology
rRNA: Ribosomal Ribonucleic acid

## Declarations

### Ethics approval and consent to participate

This study was approved by the Kenya Medical Research Institute (KEMRI) Ethical Review Board under the scientific steering committee (SSC 2776). Oral consent was obtained from each household head prior to mosquito collections.

### Consent for publication

Not applicable

### Availability of data and materials

The raw sample sequencing reads generated from this project, including those from negative (blank) and cross contamination (soil samples) controls, along with sample metadata, have been deposited in the National Center for Biotechnology Information (NCBI), Sequence Read Archive under the BioProject PRJNA672031

### Authors’ contributions

ND conceptualized and designed the study; EO facilitated and provided laboratory facilities for field work; ND, EO, MS & AL provided resources for molecular analysis; DO, MK, SO and EE performed mosquito collections, mass rearing and bioassays; DO and MS performed molecular analysis and sequencing; ND, EMN & EO supervised the work; DO and ND performed the data analysis and drafted the manuscript; all authors reviewed and approved the final version of the manuscript.

## Acknowledgements

We acknowledge and thank the homeowners in Tulukuyi village for access to their homes during mosquito collections, without which this study would not have been possible; Evans Olang and Duncan Omondi from Kenya Medical Research Institute, for assistance during mosquito collections; Maurice Ombok from Kenya Medical Research Institute for the map of Tulukuyi, and the Biotechnology Core Facility at the US Centers for Disease Control and Prevention for sequencing our samples. We are grateful to the KEMRI Director General for the permission to publish this work.

