## Supplementary material for "Western Kenyan *Anopheles gambiae s.s.* showing intense permethrin resistance harbor distinct microbiota": Suppl. 1

Suppl. 1: Summary statistics of raw sequencing outputs and resulting data following quality control and filtering

| Sample type | Resistance status (insecticide concentration) | Samples sequenced | Raw reads | Reads post QC and filtering (range) |
| --- | --- | --- | --- | --- |
| F_1_ adult | Resistant (5x) | 50 | 1,319,437 | 27,002 (3-55,795) ^a^ |
|  | Susceptible (5x) | 50 | 2,999,628 | 88,228 (5-2,697) ^a^ |
|  | **Total** | **100** | **4,319,065** |  |
| Controls | NoTemplate_Extraction Control | 4 | 305 | N/A |
|  | NoTemplate_PCRControl | 4 | 0 | N/A |
|  | SoilSample_Cross_contamination_Control | 2 | 31 | N/A |
|  | Washwater_Negative Control | 11 | 4,890 | N/A |
|  | **Total** | **21** | **5,226** |  |

QC: Quality control

^a^ Following sequencing data quality control and subsequent removal of features associated with controls and those with frequency < 100 (these were largely associated with host mitochondrial 16S rRNA), 36 susceptible and 39 resistant samples remained and were used for downstream analysis
