## Supplementary material for "Western Kenyan *Anopheles gambiae s.s.* showing intense permethrin resistance harbor distinct microbiota": Suppl. 2

Suppl. 2: Shannon alpha diversity index rarefaction curves

The number of ASVs from the microbiota of individual permethrin resistant (n = 39) and susceptible (n = 36) *An. gambiae s. s.*, along with the depth at which rarefaction was performed (100 ASVs per sample). The rarefaction plots show average Shannon diversity index and range (boxplots: minimum, median and mean) for 10 iterations of Shannon diversity analysis. The rarefaction plots plateau before the depth of rarefaction, indicating that increasing sequencing depth resulted in no/negligible change in Shannon indices.

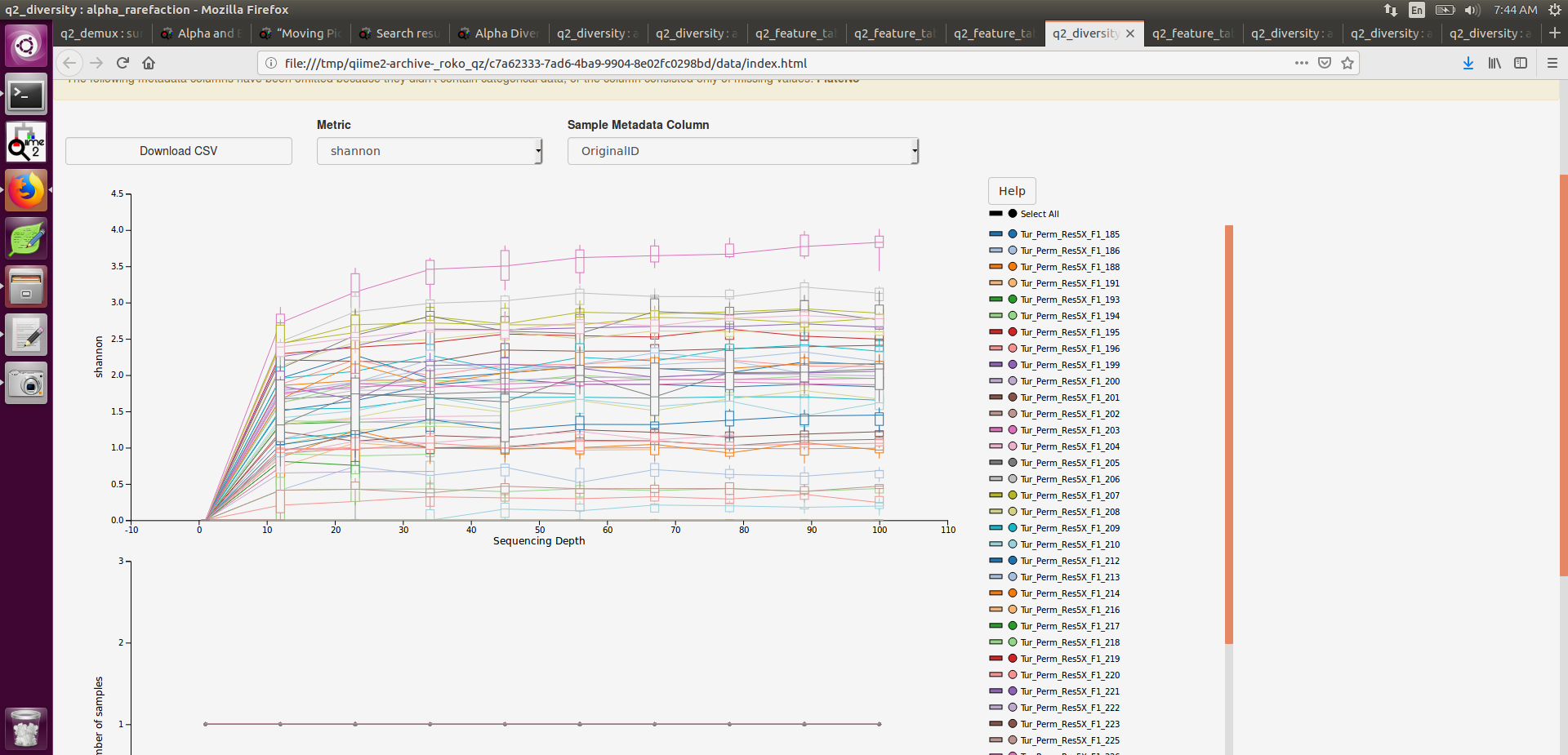
