## Supplementary material for "Western Kenyan *Anopheles gambiae s.s.* showing intense permethrin resistance harbor distinct microbiota": Suppl. 3

**Suppl. 3 : Comparison of Bray curtis dissimillarity indices between the microbiota of permethrin resistant and susceptible individuals using rarefied and unrarefeid data.**

|  | No. of mosquitoes | pseudo-F | p-value | q-value |
| --- | --- | --- | --- | --- |
| Rarefied | 42 | 1.45 | 0.02 | 0.02 |
| Unrarefied | 75 | 2.33 | 0.001 | 0.001 |

Rarefaction depth was set to 100 ASVs per sample and 33 samples that did not meet this criteria were excluded from the analysis. Comparisons were performed using 999 permutations of PERMANOVA tests with Benjamini-Hochberg FDR correction (q-value), and dignificance was set to q<0.05. There were significant differences between the microbiota of resistant and suscetible mosquitoes using either rarefied or unrarefied data. Ordinations outputs of the latter are further presented (Figure 2)
