## Supplementary material for "Western Kenyan *Anopheles gambiae s.s.* showing intense permethrin resistance harbor distinct microbiota": Suppl. 4

Suppl. 4: Shannon diversity indices showed no significant difference in diversity of bacterial taxa between individual resistant (n = 39) and susceptible (n = 36) *An. gambiae s. s.* (H= 0.45, *p*= 0.50). Comparisons were performed using Kruskal-Wallis tests (H) with Benjamini–Hochberg FDR correction (q-value). Significance was determined at q<0.05

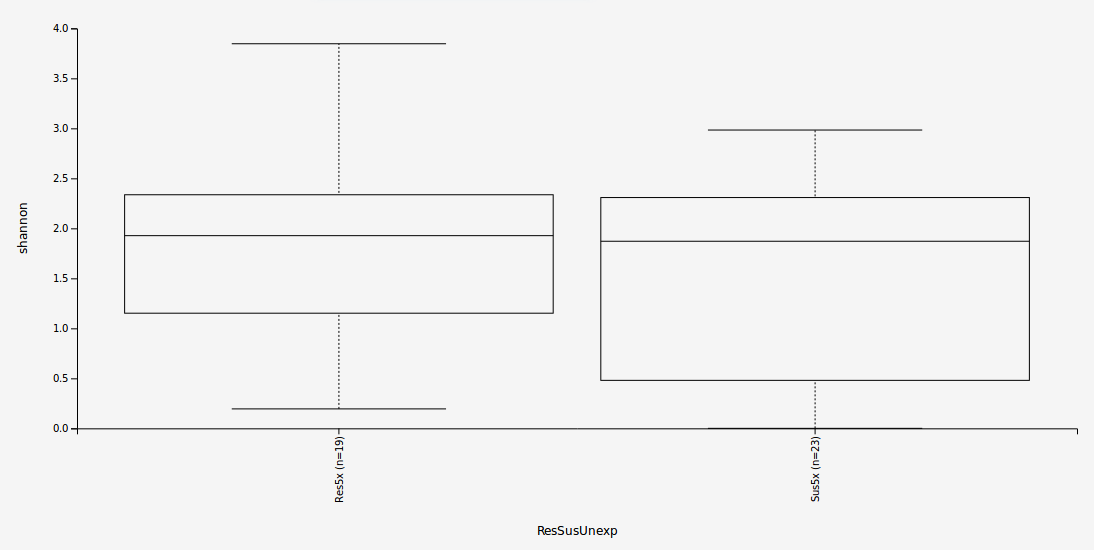
